# A population readout of extrastriate activity reveals biased and smoothed temporal representations across saccades

**DOI:** 10.64898/2026.06.16.732385

**Authors:** Adeleh Poursadegh, Maryam Zekri, Geyu Weng, Amir Akbarian, Kelsey Clark, Hossein Rabbani, Behrad Noudoost, Neda Nategh

## Abstract

Perception of visual time is transiently distorted around saccadic eye movements, yet the neuronal mechanisms constructing perisaccadic representations of time remain unclear. Here, we investigate how perisaccadic temporal information is encoded and read out in extrastriate cortex, by combining electrophysiological recordings in V4 and MT neuronal population in macaque monkeys under high spatiotemporal resolution visual stimulation, and a statistical modeling framework capturing perisaccadic response modulations at single-trial precision. Our analyses show that perisaccadic neuronal responses systematically shift the temporal representation of presaccadic stimuli within receptive fields–biasing them toward earlier times, and also reduce temporal sensitivity for presaccadic stimuli–impairing discrimination of stimulus onset times. Model-based readout using time-varying spatiotemporal sensitivity maps in neuronal ensembles enables quantitative characterization of these effects and identifies their specific neuronal response components at millisecond resolution. In silico manipulations further demonstrate a causal role of representational bias in reducing temporal sensitivity. These findings suggest that extrastriate cortex implements an active encoding strategy to stabilize presaccadic temporal information by favoring the most recent reliable input, revealing a fundamental tradeoff between temporal precision and robustness that supports a continuous visual percept across saccades. This finding also establishes a general role for extrastriate populations in constructing the perception of visual time.

## Introduction

Time perception, the cognitive capacity to quantify duration, is a ubiquitous function essential for motor control and decision-making in humans and nonhuman primates (NHPs)^1–5^. Behavioral studies demonstrate that subjective time is susceptible to contextual factors, attention, and predictability^6,7^. Neural investigations suggest that temporal processing is implemented across widely distributed networks. In NHPs, areas such as the Lateral Intraparietal area (LIP) exhibit ramping activity encoding elapsed time and reflecting evolving temporal judgments and spatial attention allocation^8,9^. Additionally, studies in humans using transcranial magnetic stimulation confirm a causal involvement of visual cortices V1 and the middle temporal area (MT) in both the encoding and working memory maintenance of temporal information in the hundreds of milliseconds range^10^.

Actions, such as movement preparation or performing manual tasks, have been shown to significantly influence our subjective perception of time^11,12^. Saccadic eye movements, which are rapid shifts in gaze, are a prime example of actions that induce temporal distortions. These quick ballistic movements are crucial for directing our gaze to areas of interest, allowing us to focus on different parts of a scene multiple times per second^13–15^. Studies have shown that visual perception of brief stimuli, presented close to the time of a saccade, can be distorted in several ways^16–18^. One well-documented distortion is mislocalization, a change in the perceived location of a perisaccadic stimulus^19–25^. Temporal distortions also occur around saccades, including phenomena of order inversion^26–28^, temporal compression (underestimation of stimulus duration or interval)^16,18,29–34^, and chronostasis (overestimation of stimulus duration or interval)^35–40^.

These perceptual distortions are accompanied by alterations in visual responses around the time of saccades^41^. Studies showed that receptive fields (RFs) actively shift perisaccadically^42–44^, a possible neural substrate for mislocalization. Furthermore, visual sensitivity decreases across much of the visual field^13,15,45^, contrasted by an increases at the target of the saccade^24,46,47^. At the same time, response latencies in visual neurons are reduced in areas MT and MST^30,31^, whereas V4 demonstrates no significant latency alterations. However, within V4, the ratio of response amplitudes to sequential stimuli undergoes perisaccadic modulation^48^. Rigorously linking observed perisaccadic changes in neural responses to their perceptual counterparts, however, is challenging, in part due to the rapid (millisecond) timescale on which neural sensitivity is changing and the difficulty in fully experimentally characterizing these dynamic modulations.

To examine the link between perisaccadic visual responses and time perception, our research utilizes a model-based approach. We use a time-varying extension of the classical generalized linear model (GLM)^49–52^ framework capable of capturing the fast spatiotemporal dynamics of neural sensitivity around the time of saccades^53^. This computational method is specifically designed to precisely characterize the dynamics of visual information encoding on the millisecond timescale, enabling temporally-precise tracing of perisaccadic sensitivity changes from neural data. Classically, the perisaccadic stimulus-response relationship has been studied using limited variations in the times and locations of visual stimuli preselected by the experimenters. However, the prevalence of reported effects often varies across studies. These discrepancies likely arise from experimental paradigms limited by coarse sampling of space or time, which fail to capture the continuous dynamics of the perisaccadic transition. To alleviate the subjectivity of this approach, we present white noise probes with high temporal precision and covering a wide range of space, and use statistical frameworks to provide an unbiased assessment of the computations underlying the neuron’s response generation. The model, fit to electrophysiological data, provides a mathematical description of the computations that the neuron applies on a stimulus to generate its response, often referred to as the neuron’s processing “kernels”, which characterize the evolving sensitivity of neurons on the fast timescale surrounding saccades. This model has been previously deployed to study the mechanism of transaccadic integration^54^ and perisaccadic mislocalization^25^. Applying this model-based approach, we aim to characterize the contributions of perisaccadic neural changes in extrastriate visual areas to temporal perception using an experimentally tractable amount of data.

This study investigated how the temporal characteristics of neural activity change around saccades. We focused on two brain regions in rhesus macaque monkeys, the MT cortex and area V4, and measured the spiking activity of neurons while the monkeys performed a visually guided saccade task with visual probes stimulation. To quantify neural sensitivity and its perisaccadic modulations, we fit the GLM-based model described above^53,55^ to the recorded neuronal responses. We used the spatiotemporal kernels constructed by the model for each group of neurons (’ensemble’), and examined how their temporal properties evolved across saccades. We found that near saccade onset, the ensemble spatiotemporal kernels exhibited higher similarity between stimulus onset times, indicating a reduction in temporal discriminability consistent with filtering out high-frequency, unreliable temporal fluctuations. Further analysis revealed a systematic change in the ensemble spatiotemporal kernels such that presaccadic representations increasingly resembled those from earlier time points, introducing a systematic bias in perceived time of presaccadic stimuli toward earlier times. The observed temporal bias suggest that stimuli presented just before a saccade may be perceived as stimuli that were available earlier, thereby extending the most recent stable presaccadic information up to just before the saccade onset. Importantly, our model allows for causal interrogation of these effects through in silico manipulations. By selectively altering components of the estimated spatiotemporal kernels, we demonstrate that the observed temporal bias leads to a reduction in temporal sensitivity, defined as the ability to discriminate between stimulus onset times. This reduction in sensitivity suggests that perisaccadic distortions are not simply a consequence of noisy or unreliable visual responses driven by rapid motor signals, but rather reflect a structured change in encoding that trades temporal precision for perceptual continuity. Overall, these findings imply that extrastriate areas actively preserve temporal continuity in visual representations, compensating for disruptions and uncertainty in temporal information caused by saccades.

## Results

### Characterizing the spatiotemporal sensitivity of extrastriate neurons across saccades

To investigate temporal processing across saccadic eye movements, we conducted electrophysiological recordings of single-unit activity from neurons in the extrastriate visual areas V4 and MT of macaque monkeys. Across 108 recording sessions, we obtained data from 300 MT and 147 V4 neurons in four adult male rhesus macaques. Neural activity was recorded using 16-channel linear array electrodes, and spikes were sorted offline. The monkeys performed a visually guided saccade task designed to probe perisaccadic visual responses (Fig. 1a). Before, during, and after a visually guided saccade, brief visual probe stimuli (white squares, 0.5×0.5 degrees of visual angle (dva); 7 ms duration) were presented within a 9×9 grid of spatial locations covering the fixation point (FP), saccade target (ST), and the neurons’ estimated RFs both before and after the saccade. The probe grid dimensions and spacing were individually adjusted in each recording session to ensure adequate coverage of the neuron’s pre- and post-saccadic RFs and surrounding visual space. Every 7ms, location of the probe was chosen randomly from the 9×9 grid of possible locations. Figure 1b shows the grid arrangement and RF locations for neurons in an example ensemble, and the firing rate of three sample neurons to the same probe in the fixation and perisaccadic period is shown in Fig. 1c. This high-density visual stimulation protocol allowed us to sample neuronal responses with fine spatial and temporal precision, thereby capturing the rapid changes in visual sensitivity that occur around the time of saccades.

**Fig. 1:**
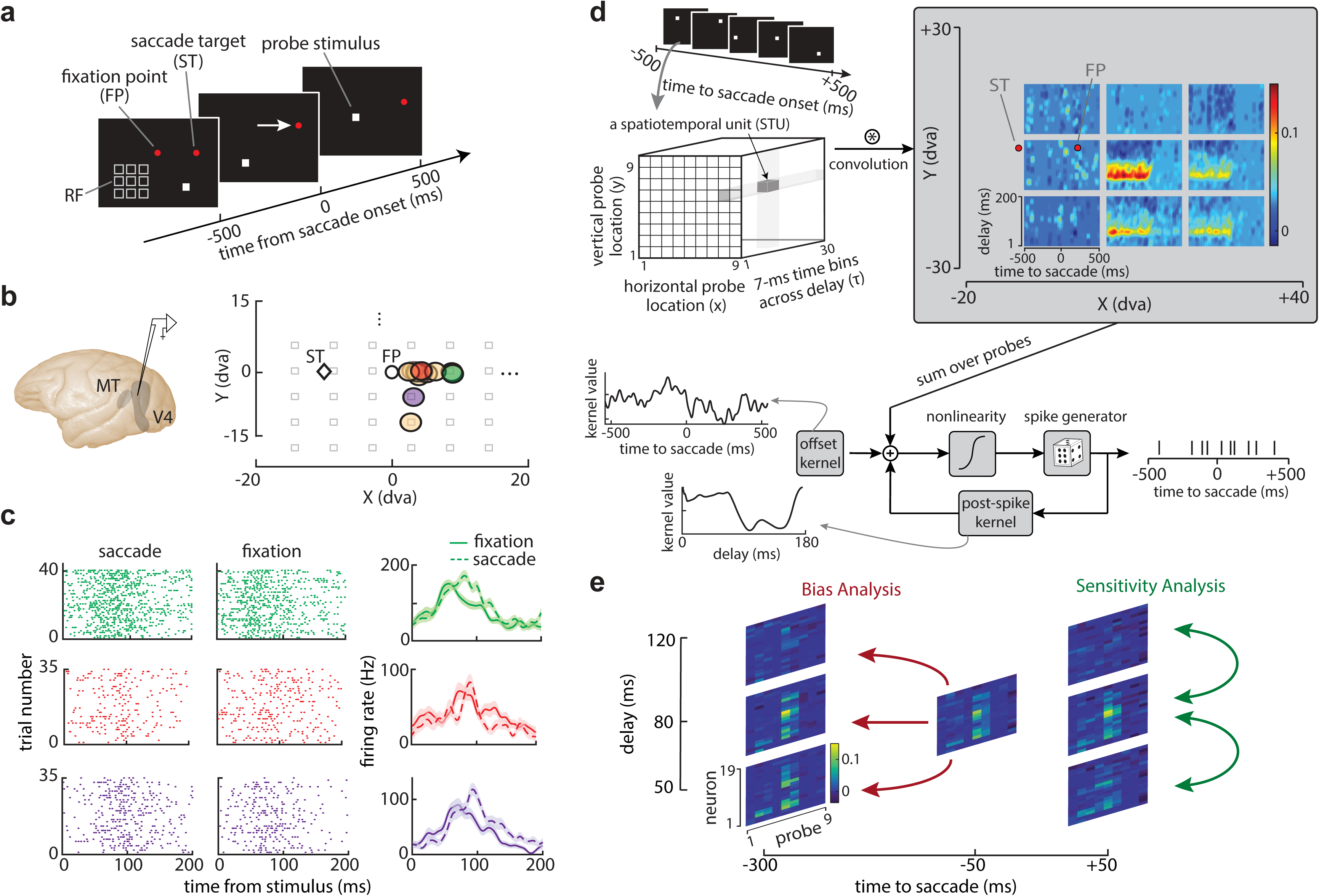
Illustration of experimental design and model-based analysis of time perception during saccades. **a.** Visually guided saccade task with pseudo-random visual probes. Each trial began with fixation on a central fixation point (FP) for 700–1100 ms. Following the disappearance of the FP, the monkeys executed a saccade toward a saccade target (ST) and maintained fixation there for an additional 560–750 ms. Brief (7ms) visual probes appeared on the screen where its location was randomly chosen from a 9×9 grid of possible locations throughout the task; only the 9 probe positions nearest to the RF center are shown here (white outlines). **b.** Left: Recordings were made from V4 or MT neurons. Right: Example spacing and positioning of probe grid (grey outlines), FP and ST locations, and receptive field (RF) center locations for multiple neurons from a representative ensemble (n=19 in this ensemble). **c.** Spike raster plots and average firing rates of three example neurons from the ensemble in (b) in response to RF center probes appearing during fixation (solid lines) or perisaccadically (dashed lines). Colors correspond to the plots of RF center positions in (b); shading indicates standard error of the response across probe presentations. **d.** Sparse variable generalized linear model (SVGLM) used to model single-neuron spiking activity. Top left: Each spatiotemporal unit (STU) corresponds to a particular probe location (x,y), time to saccade, and delay between probe and response. Top right: STU weights are fitted to a neuron’s responses, and these weighted STUs comprise a neuron’s fitted kernels. Fitted kernels are shown across time and delay values, and nine probe locations for an example neuron. Bottom: Schematic of SVGLM model structure. Stimulus inputs are convolved with kernels. The stimulus-driven component is combined with an offset term capturing baseline fluctuations and a feedback filter accounting for recent spiking history. The total input drive is transformed by a static nonlinear link function to yield an instantaneous firing rate, which serves as the intensity parameter of a conditional Poisson process generating model-predicted spike trains. Examples of fitted offset and post-spike kernels are shown. **e.** Example population kernel matrices summarizing ensemble sensitivity within the RF, shown as a function of time relative to saccade onset and stimulus delay for the representative neuronal ensemble in (b). Sensitivity analysis involved analyzing correlations between these matrices at different delays for each time to saccade. In the bias analysis, correlation of the ensemble kernel at a certain time relative to the saccade was calculated with the ensemble kernel at different delays during fixation time window.

In order to investigate the mechanism of perisaccadic temporal distortions, we leveraged a statistical encoding model that quantitatively characterizes the neuron’s input-output relationship as varying across saccades (Fig. 1d)^53^. This model captures the neuron’s sensitivity map with high temporal precision throughout the eye movement task at any time relative to saccade onset, delay between the stimulus presentation and the response time, and location. First, we decompose this four-dimensional space defined across various times and locations into discrete bins of ∼3-6 dva and 7 ms time bins (resolution of probes), to represent this space as a collection of discrete spatiotemporal sensitivity units (STUs; Fig. 1d). These STUs are used to quantify each neuron’s spatiotemporal sensitivity. In our experiment, for a 200 ms delay period over 1000 ms of response times relative to saccade onset, this space decomposition generates ∼10^7^ STUs. Using an unbiased statistical dimensionality reduction procedure, only those STUs that significantly contribute to the stimulus-response relationship are selected, resulting in discarding ∼99.9% of STUs and leaving ∼10^4^ STUs per neuron for the model estimation stage. The contribution (weight) of these selected STUs is then determined by fitting an encoding model to single-trial spiking data using point process maximum likelihood estimation. The weighted combination of these STUs defines the neuron’s stimulus kernels *k_x_*_,*y*_(*t*, *τ*), which represent the neuron’s dynamic sensitivity across different delays relative to the stimulus onset (*τ*), and locations (*x*, *y*), for any specific time relative to the saccade onset (*t*).

To fit the model to neuron’s responses and estimate STU weights and the resulting kernels, we leveraged a recent computational modeling framework, the sparse-variable GLM^53,55^ (SVGLM; Fig. 1d). This framework extends the classical GLM^49–51^ to capture the fast, high-dimensional dynamics of neural spiking that accompany saccadic eye movements using time-varying kernels. The model processes stimulus coordinates-both spatial and temporal- by convolving them with the time-varying spatiotemporal kernels that define the neuron’s specific sensitivity profile. This filtered input is integrated with two additional components: an offset kernel that accounts for baseline fluctuations synchronized with saccade initiation, and a feedback signal derived from a post-spike filter to incorporate previous firing patterns. This summed activity is transformed via a sigmoidal nonlinearity capturing the response nonlinearities, such as threshold, amplification, and saturation. Finally, a Poisson process converts these firing rates into discrete spikes, which are subsequently fed back into the system through the post-spike history mechanism. Examples of fitted spatiotemporal kernels for a sample neuron at 9 probe locations are shown in Fig. 1d. See Supplementary Figure 1 and Methods section for more details on model fitting and performance. The SVGLM is designed to model the dynamic relationship between stimulus input and neural spiking activity at a fine temporal scale, allowing for precise quantification of how neuronal sensitivity evolves during saccades; it was previously used to study the neural correlates of trans-saccadic integration^54^ and mislocalization^25^ phenomena. In this study, we investigate the possible neural mechanisms of specific alterations in time perception during a saccade.

### A model-based readout of perisaccadic temporal sensitivity

To conduct all our analyses at the population level, we grouped neurons into 15 ensembles, each containing at least 10 neurons recorded under similar RF, ST, and probe grid configurations; of these, 6 were V4 ensembles and 9 were MT ensembles. For each ensemble, the RF area is defined as the center of mass of the RF center of all ensemble neurons plus the 8 surrounding probes. The distribution of all neurons’ RF centers across all ensembles is shown in Supplementary Figure 2. For each particular time and delay, we constructed a population kernel matrix consisting of the kernel values at that specific time and delay, across RF probe locations and all neurons in an ensemble (examples in Fig. 1e). These population kernel matrices for probes in the RF area are the basis for subsequent analyses. We first quantified ensembles’ temporal sensitivity, i.e. the ability to discriminate between visual events occurring at nearby time points. We reasoned that greater similarity between ensemble kernels at different delays relative to stimulus onset would result in reduced temporal sensitivity. To quantify this, we computed the correlation between the population kernel matrices at a specific delay and other delay bins at each time to saccade onset (Fig. 1e, see Methods section). The correlation was measured for each delay bin and repeated at different times to saccade spanning the pre-, peri-, and post-saccadic windows; thus, at each time to saccade onset (10 ms bin), we have a matrix of correlation values with each element corresponding to the correlation in population kernels between delays of 0-200 ms (5 ms bin). Examples of these correlation maps at different times to saccade are shown for a sample ensemble in Fig. 2a. For each delay bin, we took the correlation values from that delay to each later delay and fitted a sigmoidal function (example in Fig. 2b), which can be thought of as the sensitivity of each delay bin to probes in other bins at each time to saccade. To measure temporal sensitivity, we defined an index called ‘time difference’, which is inversely related to temporal sensitivity. Time difference at each time to saccade onset and delay was defined as the difference between the measured delay at the inflection point of the fitted sigmoidal function and the actual delay (the delay from which correlation values were taken). These time difference values were calculated across time and delay for each ensemble. Figure 2c shows the time difference map for a sample ensemble, and Fig. 2d shows the average normalized time difference map across 15 ensembles. Figure 2e-f shows the example and the ensemble-averaged time difference for delay 40:60 ms (baseline corrected, see Methods). We observed an increase in time difference of 3.21±1.94, *p*=1.90×10^-4^ from -45:0 ms relative to saccade onset compared to during fixation (-300:-150 ms relative to saccade onset), indicating reduced temporal sensitivity prior to saccade onset.

**Fig. 2:**
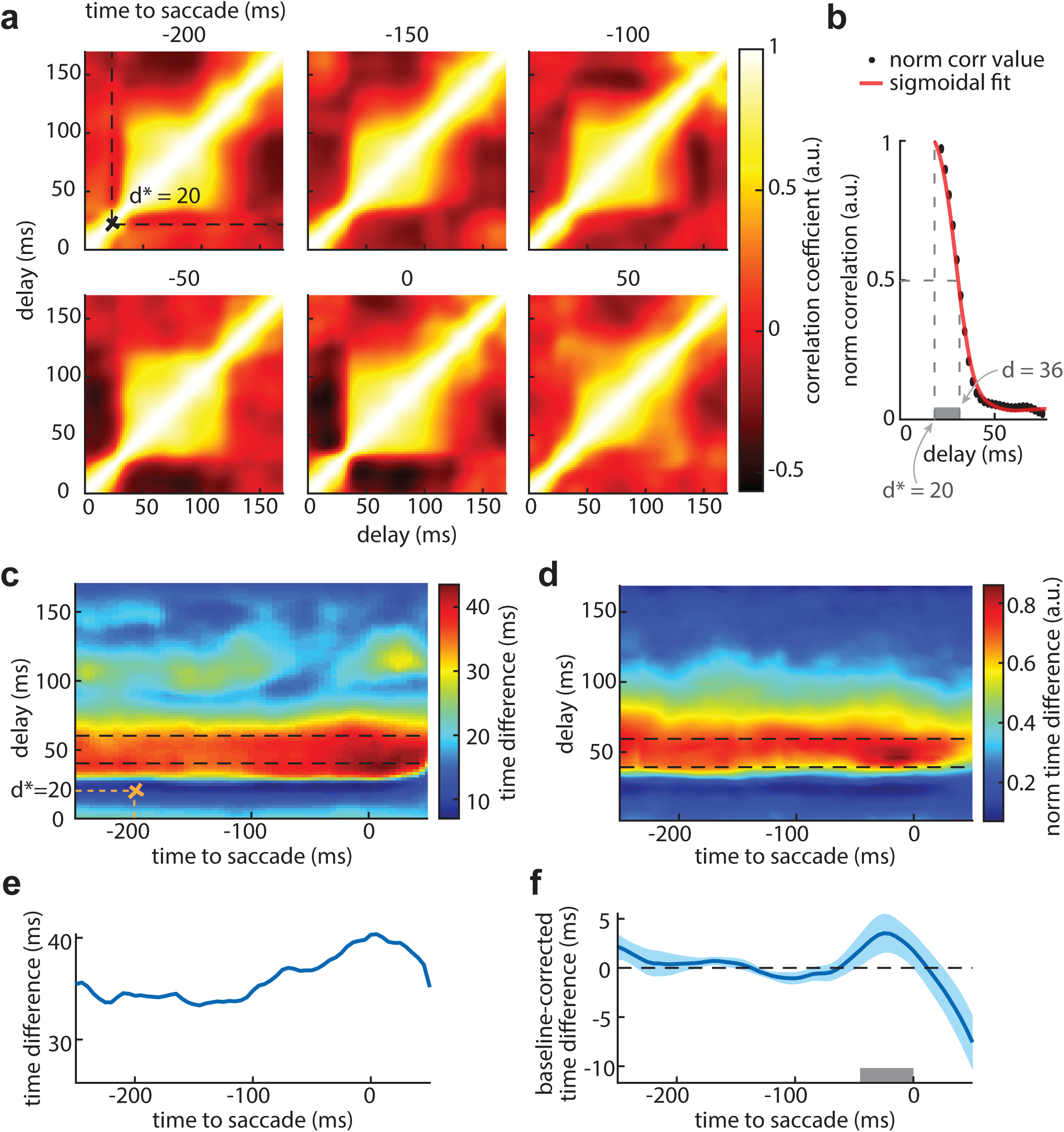
Model-based quantification of temporal sensitivity. **a.** Example correlation maps for a representative neuronal ensemble, for various times to saccade. For each time relative to saccade onset, correlations between population kernels at different delays were computed. Dashed lines in upper left plot indicate the correlation values which are used for the sigmoidal fit shown in (b). **b.** Example sigmoidal fit of correlation values across delays, for an actual perisaccadic delay of 20 ms at a time of −200 ms relative to saccade onset. The difference between the delay at the inflection point of the fitted curve (correlation value of .5; here, 36 ms) and the true probe delay (20 ms) was taken as a measure of temporal sensitivity. **c.** Time difference values for the sample ensemble, mapped across delay and time to saccade. **d.** Normalized average time difference map across 15 neuronal ensembles. **e.** Average time difference map for the sample ensemble, computed over delays of 40–60 ms. **f.** Baseline-corrected mean time difference averaged over delays of 40–60 ms across 15 ensembles (n = 447 neurons). Values are shown as mean ± SEM across ensembles. Time difference is significantly elevated from −45 to 0 ms (gray bar; *p*=1.90×10^-4^).

### Perisaccadic temporal sensitivity measured using neuronal responses

To validate the model-derived predictions using neurophysiological activity, we applied the same analysis directly to the recorded neuronal probe-aligned spiking data. We used the actual firing rate of the ensemble of neurons in response to probe stimuli inside the RF area to compute the correlation between different delay bins at each stimulus time relative to saccade onset, as shown for an example ensemble at select timepoints in Fig. 3a. We then fitted a sigmoidal function to each normalized cross-section of these maps, and time difference was calculated as the difference between the point of equality of the fitted curve and the actual delay. As with the model data, we then computed the time difference across delay and stimulus time relative to saccade onset for probes presented inside the RF area. The resultant time difference map is shown for a sample ensemble in Fig. 3b. The normalized average result across all 15 ensemble of neurons is shown in Fig. 3c. The time differences for delays of 40:60 ms are shown for the example ensemble in Fig. 3d, and across ensembles in Fig. 3e (baseline-corrected, see Methods). The neuronal results closely paralleled those predicted by the SVGLM ensemble kernels, showing a clear increase in time difference (4.61±2.27, *p*=5.24×10^-5^) measures around time window of -100:-40 ms from stimulus onset (compared to the fixation window of -700:-400 ms), indicating a perisaccadic decline in temporal sensitivity. Note that for the kernel analysis (Fig. 2), time is defined as referenced relative to the onset of a saccade, while response data (Fig. 3) are based on the time of stimulus onset. Importantly, this time relative to stimulus onset aligns with the response time window (-45:0 ms, as described in the previous section) used to assess temporal sensitivity, reflecting the typical 40–60 ms response latency of V4 and MT areas.

**Fig. 3:**
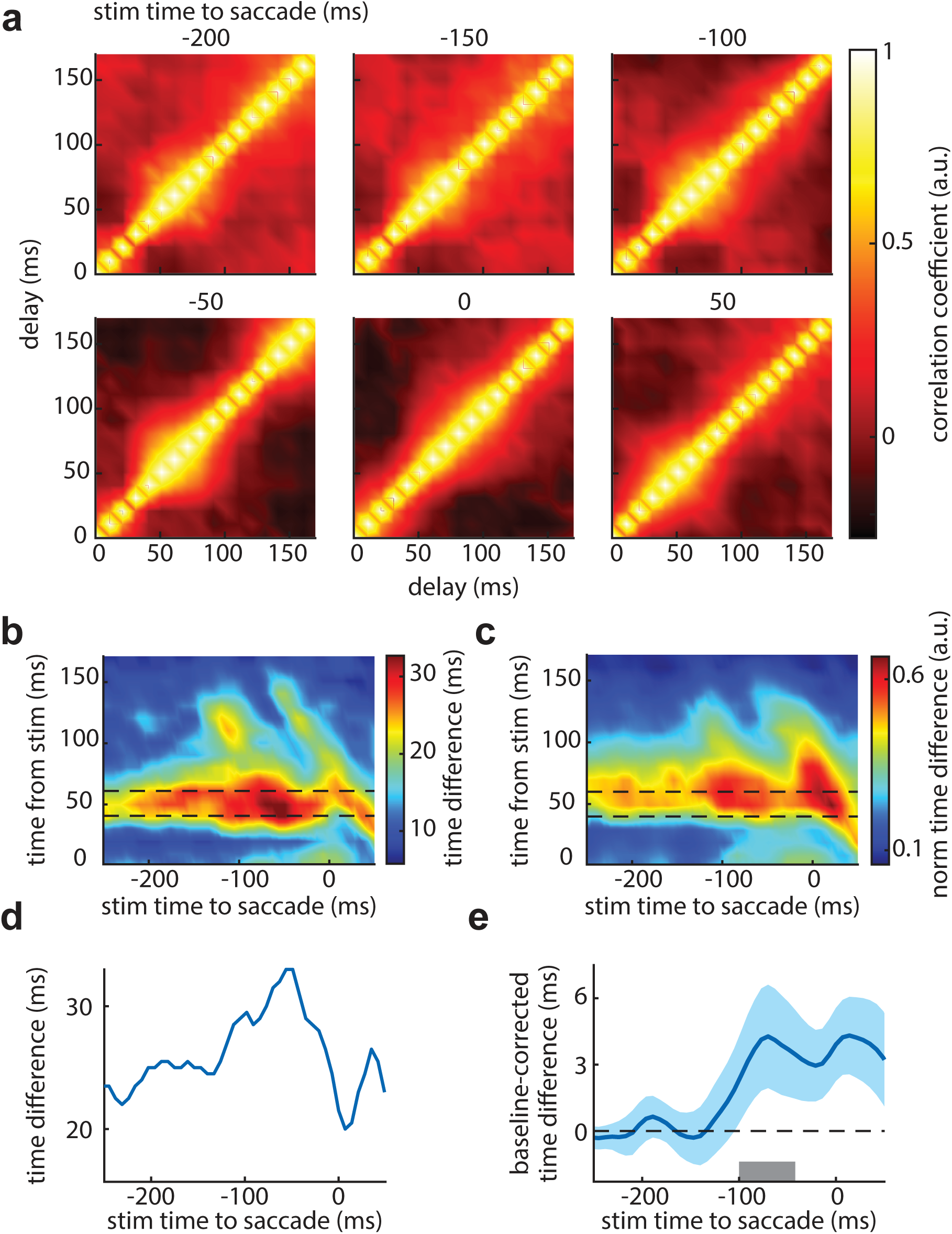
Ensemble-based quantification of temporal sensitivity from neuronal responses. **a.** Example correlation maps for a representative neuronal ensemble, time of stimulus relative to saccade onset indicated at top of each panel. For each time, correlations between firing rates at different delays relative to stimulus onset were computed. **b.** Corresponding time-difference map for this ensemble, across delay and time of stimulus to saccade. **c.** Normalized average time-difference map across all 15 neuronal ensembles. **d.** Average time-difference map for the sample ensemble, computed over delays of 40–60 ms. **e.** Baseline-corrected mean time difference averaged over delays of 40–60 ms across ensembles (n = 447 neurons). Values are shown as mean ± SEM across ensembles. Time difference significantly increased from −100 to −40 ms (gray bar; *p*=5.24×10^-5^).

We hypothesize that the observed reduction in temporal sensitivity could arise from one of two potential mechanisms (schematically illustrated in Fig. 4a, b). First, it could result from increased noise (high frequency fluctuations) in the temporal representation during the perisaccadic period, reflecting a degradation of the precision with which stimulus timing is encoded. Alternatively, it could reflect a systematic temporal bias, whereby neural representation of perisaccadic stimuli at each timepoint is temporally shifted toward certain timestamps, which may lead increased representational similarity and thereby a reduction in the ability of the response to discriminate between different timepoints. To differentiate between these possibilities, we conducted a second set of analyses aimed at characterizing the temporal mapping properties of neural population activity during the perisaccadic interval.

**Fig. 4:**
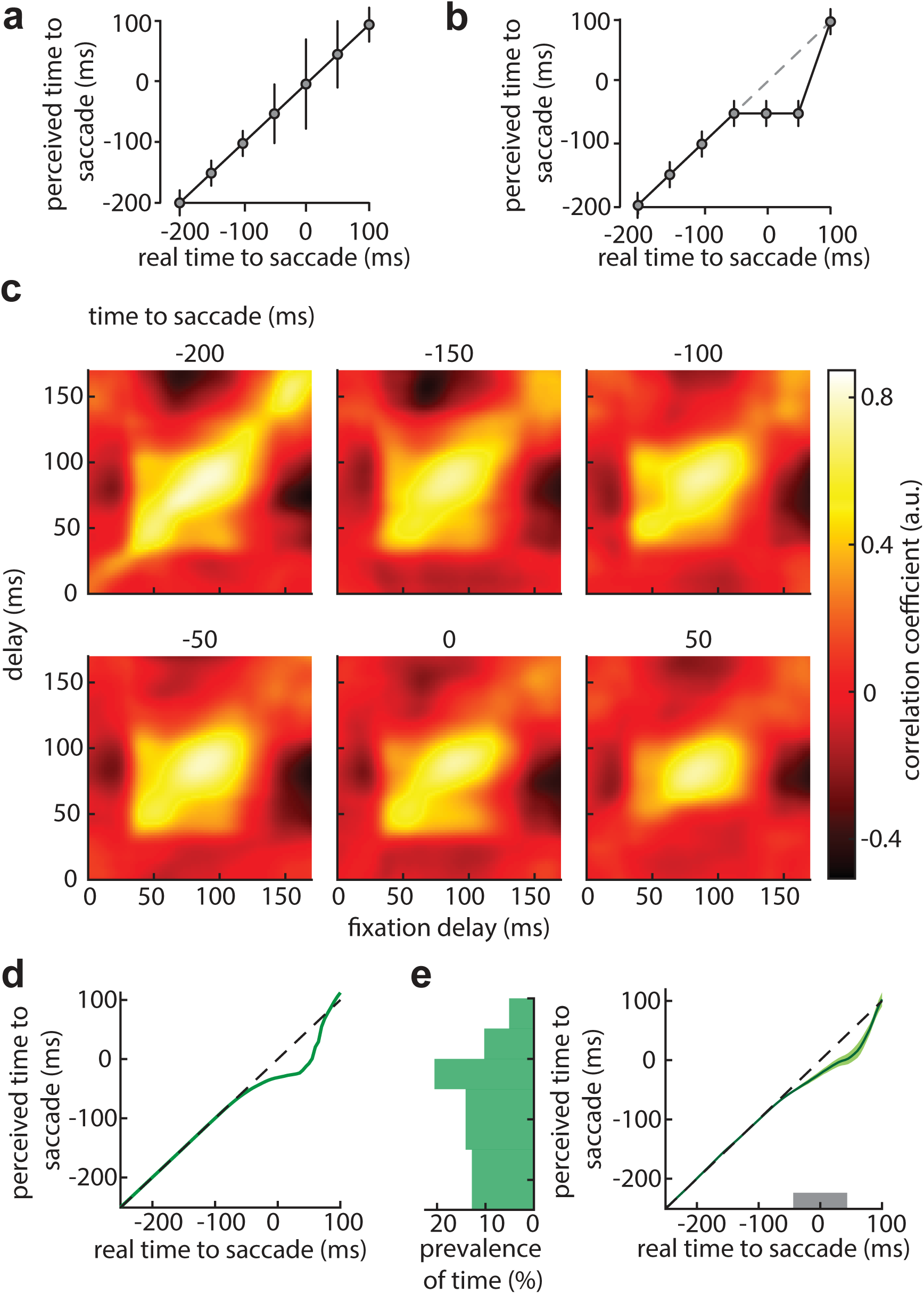
Model-based quantification of temporal bias from population kernels. **a.** Schematic of relationship between real and perceived time with increased noise. **b.** Schematic of relationship between real and perceived time with systematic perisaccadic bias. **c.** Example correlation maps for a representative neuronal ensemble, at various times to saccade (indicated on top of each panel). Plots show the correlation between population kernels at delays from the fixation period and those for delays from the time indicated at top. **d.** Perceived versus real time for this ensemble, obtained by averaging across time points within a 300-ms sliding window. The correlation of each time point’s map with the central map was used as a weighting factor for its contribution to perceived time. **e.** Average temporal bias across 15 neuronal ensembles. Left histogram shows the distribution of perceived time to saccade. Shaded regions denote the SEM across ensembles (gray bar; *p*=5.50×10^-4^).

### Model-predicted bias in the representation of time information across saccades

Assuming that temporal encoding is incorporated into ensemble kernels during fixation, we investigated how this representation could give rise to perceived time around saccade onset. To assess how temporal representations are altered around the time of saccades, we examined the correlations between perisaccadic and fixation kernels across delays. These correlations served as a measure of how neural representations at different time points were mapped onto one another. We measured the similarity between kernels at each time relative to saccade onset and those in the fixation window by computing the correlation between the ensemble kernel at a specific delay and time and the ensemble kernel at multiple delay bins in the fixation period (-500:-300 ms from saccade onset) (Fig. 1d). The correlation was measured for each delay bin across time to saccade onset. Thus, at each time to saccade onset, we have a matrix of correlation values, each element corresponding to the correlation between delays of 0-200 ms (5ms bin) and those in the fixation window. Examples of these correlation maps are shown for a sample ensemble in Fig. 4c. Next, we measured the similarity between each 300 ms sliding window of these maps by computing the correlation between each map in this sliding window and its central map, meaning that we have a scalar value of correlation associated with each time to saccade onset (see Methods section). Treating these scalars as weighting factors for each bin of time in a sliding window, and then taking sum of all the time to saccade bins in the sliding window results in the perceived time to saccade. The time to saccade corresponding to the central map is considered as the real time. Figure 4d shows an example plot for real vs. perceived time for an ensemble. The average of mapping between perceived and actual time across 15 ensembles is shown in Fig. 4e. Left histogram in Fig. 4e depicts the distribution of perceived time to saccade. From these results, we observed that time windows just before saccade onset were systematically mapped to earlier times, such that the distribution of perceived time is no longer uniform perisaccadically. To measure this phenomenon, we identified the most prevalent perceived time interval, and used it to define temporal bias as the difference between the mean perceived and real time intervals. Averaging over -50:50 ms from saccade onset across all ensembles revealed a significant temporal bias of -18.54±3.68 ms (*p*=5.50×10^-4^, n=15 ensembles). This perisaccadic temporal bias suggests that the brain’s encoding of temporal information becomes distorted around saccades, effectively shifting the perceived timing of events occurring near saccade onset earlier.

### Perisaccadic temporal bias measured using neuronal responses

We next sought to confirm these model-derived findings using directly recorded neural responses. Mimicking the analysis applied to the model, we used the stimulus-aligned firing rates of the ensemble’s neurons to generate the correlation maps between perisaccadic and fixation 10 ms delay bins across time of stimulus (15 ms bins) as shown in Fig. 5a. Again, we used a 300 ms sliding window to compute the similarity between these correlation maps, which can be seen as a weighting factor for each stimulus time bin. After summing the weighted time bins, we have the perceived time associated with each real time of stimulus, as shown for a sample ensemble in Fig. 5b and averaged across all 15 ensembles in Fig. 5c. Left histogram in Fig. 5c shows the distribution of perceived stimulus time to saccade. To quantify the temporal bias, we found the most prevalent interval of perceived stimulus time over real stimulus time of -100:0 ms from saccade onset, and defined the temporal bias as the difference between the mean real and perceived stimulus time intervals. We took the average across all ensembles, and observed -17.3±4.15 ms of temporal bias in this window (*p*=5.10×10^-3^, n=15 ensembles). Consistent with model predictions, the time window relative to the stimulus aligns with the response time window (-50:50 ms, as described in the previous section), based on the typical response latency of V4 and MT areas, which confirms the presence of structured, time-dependent distortions in neural coding.

**Fig. 5:**
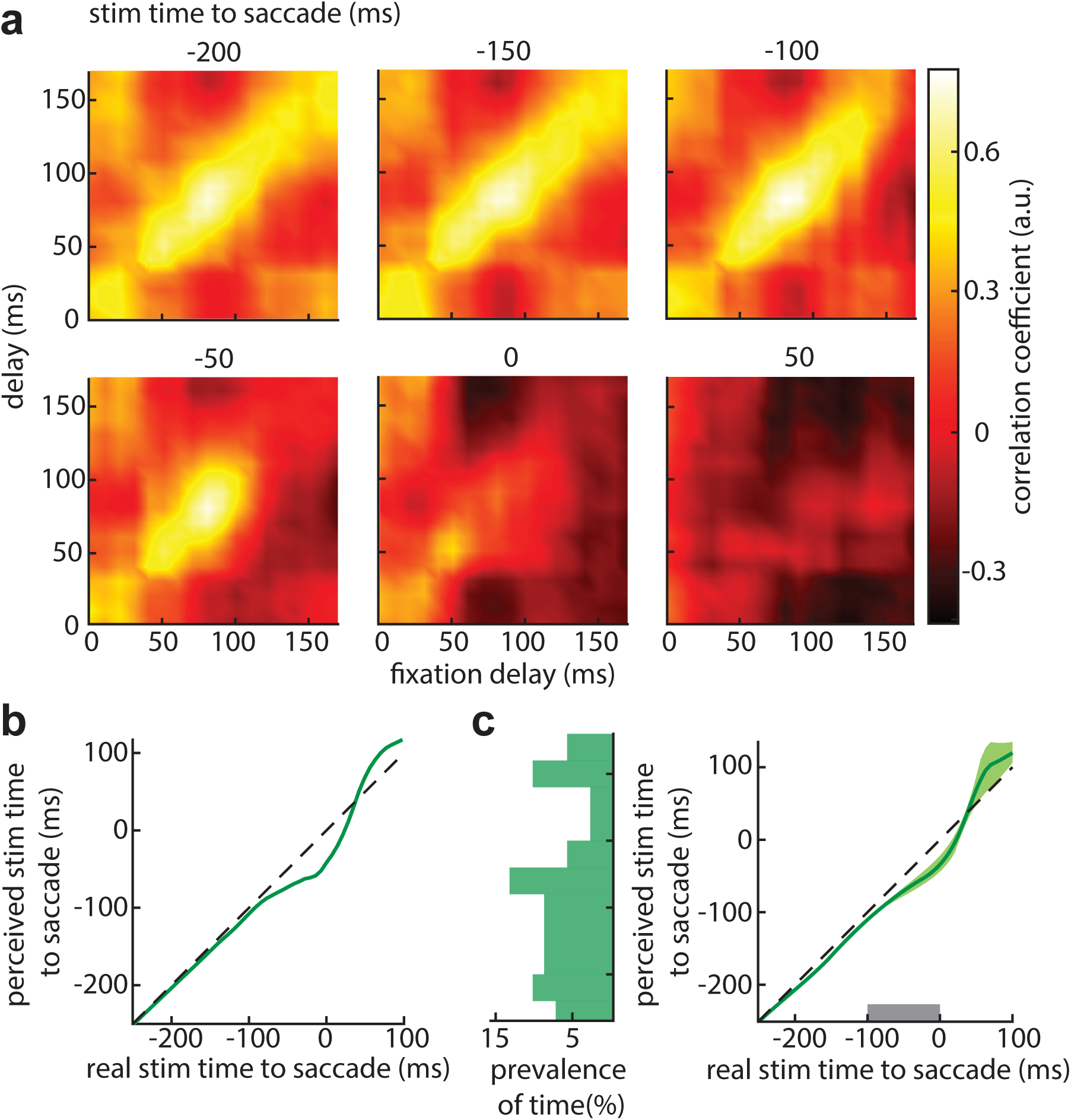
Ensemble-based quantification of temporal bias from neuronal responses. **a.** Example correlation maps at different stimulus times relative to saccades (top label) for a representative neuronal ensemble. Plots show correlations between firing rates for fixation delays with delays at time shown on top for each panel. **b.** Perceived versus real stimulus time for this ensemble, computed by averaging across time points within a 300-ms sliding window. The correlation of each time point’s map with the central map was used as a weighting factor for its contribution to perceived time. **c.** Average temporal bias across 15 neuronal ensembles. Left histogram shows the distribution of perceived stimulus time to saccade. Shaded regions denote the SEM across ensembles (gray bar; *p*=5.10×10^-3^).

### The role of representational bias in the perisaccadic readout of visual time

The SVGLM framework allows us to identify and manipulate specific components of the neural representation to determine their contribution to particular perisaccadic perceptual changes. Using this in silico approach, we systematically identified and eliminated individual STUs to characterize their role in generating specific perisaccadic temporal effects. We leveraged this feature to test the two hypotheses illustrated in Fig. 4a, b, and to identify subsets of STUs that were either responsible for generating temporal bias in the neural representation (’bias-relevant’), or associated with increased temporal variability (’noise-relevant’).

To identify bias-relevant STUs, we computed an index reflecting the difference between the stimulus kernels of each delay bin and the average kernel of delays 40:100 ms at each time to saccade. STUs were classified as bias-relevant if removing them significantly altered this index (see Methods). Figure 6a shows how the kernel weights evolve during the perisaccadic period across delay values. The prevalence map of identified bias-relevant STUs for an example ensemble of neurons is shown in Fig. 6b. Figure 6c shows the average prevalence map across all neurons, illustrating that most of the bias-relevant STUs are in the perisaccadic window. After replacing the perisaccadic weights of these bias-relevant STUs with their fixation values, we recomputed both the temporal bias and time difference over time and delay values. As expected, after replacing bias-relevant STUs, there was a significant reduction in temporal bias compared to the full model (-50:50 ms window, -6.23±3.21; *p*=2.66×10^-2^, Fig. 6d). Figure 6e shows the time difference map after removing the bias-relevant STUs, and Figure 6f demonstrates the mean time difference over delay of 40:60 ms, no longer showing a significant time difference (0.80±1.77, *p*=1.10×10^-3^), demonstrating that the same STUs drive the changes in both perisaccadic sensitivity and temporal bias. We can also test the effect of removing these bias-relevant STU modulations on the model-predicted spiking of neurons to synthetic probe sequences. We recomputed the model’s perisaccadic response after removing bias-relevant STUs via targeted kernel refitting, followed by reevaluation of temporal sensitivity and bias from the updated model-predicted spike trains. Fig. 6g confirms that the bias in temporal perception is reduced after removing bias-relevant STUs. Specifically, taking the average of difference between the most prevalent real and perceived time over stimulus time of -100:0 ms from saccade shows -12.23±1.98 ms (*p*=0.12) of temporal bias in this window. Figure 6h shows the average time difference map across all 15 ensembles of neurons. Figure 6i shows the mean baseline-corrected time difference over delay of 40:60 ms; removing bias-relevant STUs eliminated the perisaccadic increase in time difference (-100:-40 ms from saccade onset; 0.19±1.14, *p*=3.80×10^-3^), reiterating the common basis for perisaccadic changes in bias and sensitivity. In contrast, to identify noise-relevant STUs, we examined whether there is any variability of kernel weights across time to saccade, by computing the variance of the stimulus kernels of each delay bin across time to saccade for individual neurons, which may impact the resulting temporal bias read out from that ensemble (see “Methods” section). The standard variance of stimulus kernels of the fitted models across time to saccade (*t*) at each delay bin (*τ*), was used as a variability index. STUs whose removal significantly changed this variability index were labeled noise-relevant. Figure 7a illustrates the kernel weights across time to saccade onset at different delays for a sample neuron. Figure 7b, c shows the prevalence of noise-relevant STUs for a sample neuron and averaged map across all 15 neurons, respectively. Eliminating these noise-relevant STUs did not affect the temporal bias (-14.73±3.18 ms; *p*=0.25, Fig. 7d). Figure 7e depicts the time difference map computed after removing noise-relevant STUs, and Figure 7f is the average of baseline-corrected time difference map over delay of 40:60 ms, which shows that there is still an increase in time difference around -45:0 ms of saccade onset (1.93±2.10, *p*=4.82×10^-2^) even after removing the noise-relevant STUs. Thus, the observed perisaccadic bias and reductions in temporal sensitivity were not the result of noisier neural coding. We then tested the effect of removing the noise-relevant STUs on the model’s predicted spiking responses and related temporal sensitivity and bias calculations. Figure 7i shows the recomputed estimated perceived time using model-predicted response of noise-reduced model, which shows an average temporal bias of -17.97±2.16 ms (*p*=0.56) over time of -100:0 ms, which is not different from the full model results. The noise-reduced temporal sensitivity map is shown in Fig. 7h. The mean baseline-corrected time difference over delay interval of 40–60 ms is shown in Fig. 7i, with no significant change in time difference from the full model 2.73±1.71, *p*=0.11 for -100:-40 ms from saccade onset. The baseline in this plot was computed over the reference period of -350:-250 ms of stimulus time from saccade onset. Together, these results demonstrate that perisaccadic declines in temporal sensitivity are not attributable to increased variability of responses, but instead reflect systematic changes in kernel structure that shape temporal readout around saccades.

**Fig. 6:**
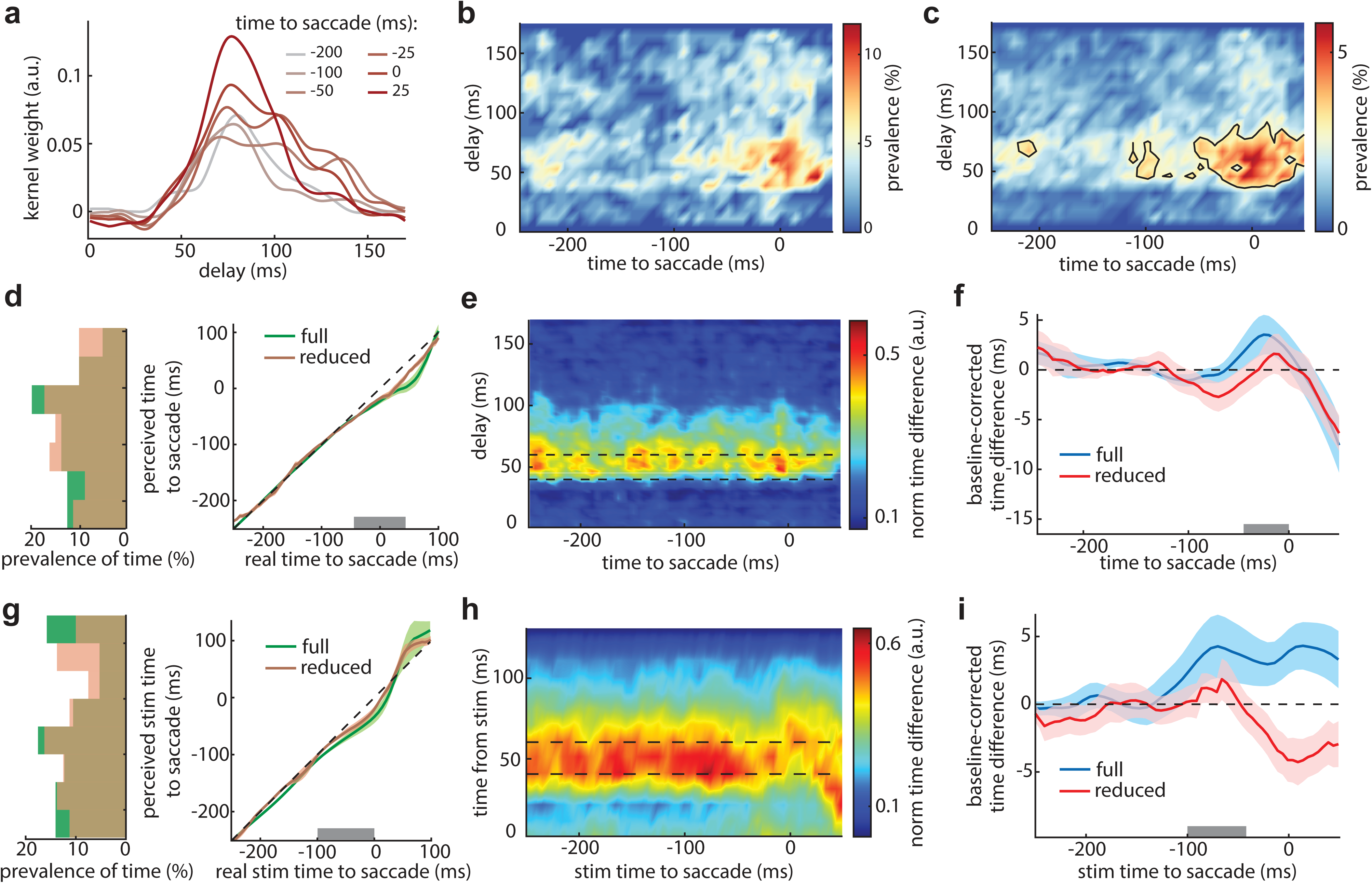
Identification and validation of bias-relevant sensitivity components. **a.** Example spatiotemporal kernel weights across delays at different times relative to saccade for a representative neuron. **b.** Prevalence of bias-relevant STUs as a function of delay and time from saccade for the example ensemble. **c.** Prevalence of bias-relevant STUs averaged across all 15 neuronal ensembles. Black outline shows area with prevalence more than half of peak for this map. **d.** Mean estimated perceived-time maps as a function of time from saccade onset for the full model (green) and the bias-reduced model (brown), averaged across ensembles, demonstrating that replacing the bias-relevant STUs eliminates temporal bias in perception. Left histogram shows marginal distribution of perceived times (gray bar; *p*=2.66×10^-2^). **e.** Mean time-difference map as a function of time from saccade for the bias-reduced model, averaged across ensembles. **f.** Baseline-corrected mean time difference averaged over probe delays of 40–60 ms for the full model (blue) and the bias-reduced model (red), in which all bias-relevant STUs were replaced with fixation kernel weights. Values represent mean ± SEM across 15 ensembles. Replacing bias-relevant STUs abolishes the perisaccadic increase in time difference in the −45 to 0 ms window following saccade onset (gray bar; *p*=1.10×10^-3^). **g.** Mean estimated perceived stimulus time across ensembles computed from bias-reduced model responses (brown) and actual neuronal responses (green). Left histogram shows marginal distribution of perceived times (gray bar; *p*=0.12). **h.** Average time-difference map across 15 neuronal ensembles computed from spike responses generated by the bias-reduced model. **i.** Baseline-corrected mean time difference averaged over probe delays of 40–60 ms for bias-reduced model responses (red) and actual neuronal responses (blue). Mean ± SEM across ensembles confirms a significant difference between full and bias-reduced model for the −100 to −40 ms window (gray bar; *p*=3.80×10^-3^).

**Fig. 7:**
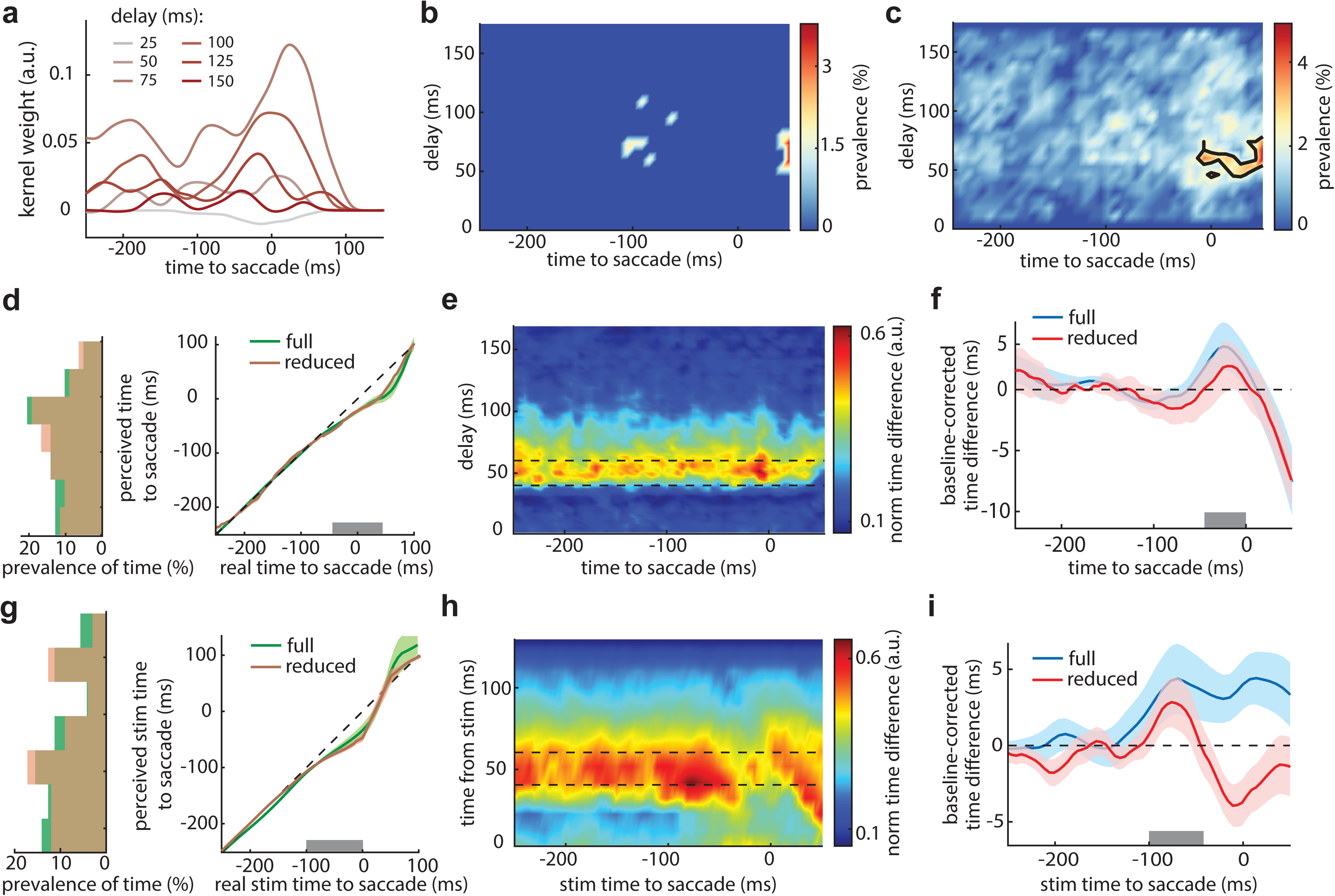
Identification and validation of noise-relevant sensitivity components. **a.** Example spatiotemporal kernel weights within the RF, shown across time relative to saccade onset at different delays for a representative neuron. **b.** Prevalence of noise-relevant STUs as a function of stimulus delay and time from saccade for the example ensemble. **c.** Prevalence of noise-relevant STUs pooled across all 15 neuronal ensembles. Black outline shows area with prevalence more than half of peak for this map. **d.** Mean estimated perceived-time maps as a function of time from saccade onset for the full model (green) and the noise-reduced model (brown), averaged across ensembles, showing that temporal bias persists after removal of noise-relevant STUs. Left histogram shows marginal distribution of perceived times (gray bar; *p*=0.25). **e.** Mean time-difference map as a function of time from saccade for the noise-reduced model, averaged across ensembles. **f.** Baseline-corrected mean time difference averaged over probe delays of 40–60 ms for the full model (blue) and the noise-reduced model (red), in which all noise-relevant STUs were replaced with fixation kernel weights. Values represent mean ± SEM across 15 ensembles. There is no significant change in the time difference in the −45 to 0 ms window for the full vs. noise-reduced models (gray bar; *p*=4.82×10^-2^). **g.** Mean estimated perceived stimulus time across ensembles computed from noise-reduced model responses (brown) and actual neuronal responses (green). Left histogram shows marginal distribution of perceived times (gray bar; *p*=0.56). **h.** Average time-difference map across 15 neuronal ensembles computed from spike responses generated by the noise-reduced model. **i.** Baseline-corrected mean time difference averaged over probe delays of 40–60 ms for noise-reduced model responses (red) and actual neuronal responses (blue). Mean ± SEM across 15 ensembles. There is no significant change in the time difference in the −100 to -40 ms window for the full vs. noise-reduced models (gray bar; *p*=0.11).

## Discussion

Saccades induce a range of distortions in the perception onset time or duration^11^, including the subjective lengthening of perceived durations (chronostasis)^35^, and the compression of temporal intervals^29^. These phenomena challenge the notion of a stable and continuous internal clock^56,57^, pointing instead to a dynamic and context-dependent construction of subjective time. In the present study, we used a combined electrophysiological and computational framework to investigate the link between perisaccadic modulations of neuronal responses in extrastriate cortex and temporal perceptions. Our results show a reduction in temporal sensitivity (the ability to discriminate between different stimulus onset times), and a bias toward perceiving stimulus onset as earlier. These findings suggest that presaccadic shifts in perceived timing play a key role in maintaining a smooth and continuous visual experience. First, they transiently hold on to the visual information available just before the saccade, effectively stabilizing the presaccadic scene until new input is processed post saccade. Second, they reduce temporal sensitivity, filtering out high-frequency and potentially unreliable temporal fluctuations. Together, these findings underscore the role of extrastriate processing in maintaining a temporally stable representation of the visual scene, despite the jitters and uncertainty inherent in saccadic eye movements.

We found that the onset of stimuli presented -100:0 ms relative to saccade onset is biased toward earlier times inside the RF area of neurons. This phenomenon suggests that presaccadic visual signals have shifted perceptual latency such that stimuli presented just before a saccade may be perceived as stimuli that were available earlier. This effect reflects a bias in perceived timing, which could lead to apparent temporal expansion, similar to chronostasis. In saccadic chronostasis, there is subjective temporal lengthening or overestimation of the duration of a visual stimulus^35,39^. It is shown that the overestimation is not proportional to overall stimulus duration^37^, and subjects typically overestimate the time they have seen the saccadic target by about 120 ms^35^. Our observed backward shift in perceived stimulus onset prior to a saccade, however, offers a potential replay-like mechanism that preserves the most recent stable presaccadic temporal representation, being temporally distinct from postsaccadic chronostasis. Rather than simply freezing the visual scene, this active process may serve to preserve the most recent stable presaccadic temporal representation and mitigate perisaccadic uncertainty. By shifting incoming signals to earlier time points, this presaccadic temporal bias establishes a temporal prior, which favors the most recent reliable presaccadic input, enabling an extrapolation of this information forward in time. This inferential strategy, supported by the temporal prior, could facilitate an active reconstruction of the presaccadic visual scene, and contribute to continuity and stability in temporal perception across saccades. In addition, we showed that during the time window of - 45:0 ms before saccade onset, the temporal sensitivity significantly declines. We hypothesize that this perisaccadic sensitivity reduction can be attributed to the previously reported effect of early saccadic suppression^53,54,58–60^ and late enhancement of response^54^, which were also observed for a similar dataset; to elaborate, the suppression of early onset times and the enhancement of late ones could result in a rise in the similarity between onset times in perisaccadic windows. It has been shown that the saccadic suppression is accompanied by a change in detection thresholds^13,61^. Reduced temporal sensitivity within neurons’ RFs, captured by perisaccadic spatiotemporal kernel modulations of extrastriate neurons, can function as a low-pass filter, preventing the relay of high frequency saccade-induced signals to downstream areas. By sacrificing temporal precision, the brain enhances informational robustness during high uncertainty. This reduction, accompanied by the observed presaccadic temporal biases, can also generate a sensory memory for presaccadic input, which may support the temporal stabilization of the presaccadic scene, despite possible variability introduced by saccade-induced input, motor planning, and attentional reallocations affecting extrastriate neurons.

One complication in any interpretation of any perisaccadic phenomena is the possible influence of attentional deployment and its effects on perception, including temporal perception. It is known that the attention shifts toward the saccade target prior to saccade execution^62^. The effect reported here is observed for the RF area, outside the focus of attention. Allocating attentional resources to the ST may limit the amount of capacity available for processing the RF, that is resources are engaged by the ST and so are unavailable for encoding the RF area. Furthermore, the role of attention and its spatial specificity in chronostasis is contested. Some studies suggest that chronostasis is a local, object-specific phenomenon, strongest at the saccade target and diminished at positions midway between the fixation point and target, implying a link to shifts in attention that precede eye movements^63^. Other research indicates that chronostasis can occur across wider visual fields, including the initial fixation location, and is not limited to the immediate saccade target region^40^, which is consistent with our findings that temporal expansion can occur in the RF region, which can be at different positions relative to saccade target.

Further research will be needed to probe the link between the effects reported in behavioral paradigms and those shown here from an electrophysiological viewpoint, including performing a more precise analysis of whether the timecourse and spatial^40^ profile of the perceptual and neurophysiological effects align. Ideally such a comparison would occur across studies in which all task parameters (saccade vectors, visual probe size and contrast, etc) are exactly matched. Future works can also explore how other brain areas can influence the perception of time. For instance, recent human studies by Zimmermann show that time compression is tightly linked to motor planning. In double-step saccades, time perception is distorted even before the second eye movement occurs, suggesting the motor plan itself –not just the movement– warps time^33,34^. Therefore, future works can explore the role of oculomotor areas that generate the saccade, such as FEF, which has also been shown to play a role in attentional and spatial modulation of V4 activity^64,65^. Also, one study has reported a potential neural mechanism for temporal compression effect based on the properties of spatially remapped future field responses of FEF neurons^66^. In conclusion, we have shown that extrastriate cortex implements a dynamic temporal encoding mechanism that shifts and smooths time representations during saccades, enabling transsaccadic temporal stability while producing systematic perceptual distortions such as temporal expansion and reduced temporal sensitivity, despite the inherent disruptions and uncertainties associated with saccadic eye movements.

## Methods

### Experimental paradigm and electrophysiological recording

All procedures complied with the NIH Guide for the Care and Use of Laboratory Animals and were approved by the Institutional Animal Care and Use Committee of the University of Utah. Four adult male rhesus macaques (Macaca mulatta) performed a visually guided saccade task. Monkeys fixated on a central point (FP) while a peripheral saccade target (ST) appeared 10–15 degrees of visual angle (dva) away. After a randomized interval (700–1100 ms), the FP disappeared, cueing the monkey to saccade to the ST. A liquid reward was delivered after maintaining fixation on the ST for 560–750 ms. In most sessions (85 out of 108), the ST location was fixed; in the remaining sessions (23 out of 108), it was randomly positioned to the left or right of the FP.

Throughout each trial, 81 task-irrelevant probe stimuli (white squares, 0.5×0.5, 7 ms duration) were presented sequentially without overlap in a pseudorandom order. Probes were arranged in a 9×9 grid scaled to cover the estimated pre- and postsaccadic receptive fields (RFs), FP, and ST. Grid dimensions varied horizontally (24–48.8 dva, mean ± SD = 40.6 ± 5.9 dva) and vertically (16–48.8 dva, mean ± SD = 39.8 ± 7.8 dva). For MT neurons, direction preference was assessed using a full-field Gabor paradigm prior to the saccade task.

Spiking activity and local field potentials (LFPs) were recorded from areas V4 and MT using 16-channel linear array electrodes (Plexon V-probes) at 32 kHz. Data were acquired using Blackrock (Central v7.0.6) and Neuralynx (Cheetah v5.7.4) systems, and spikes were sorted offline (Plexon Offline Spike Sorter, BOSS). Eye position was monitored at 2 kHz using an EyeLink 1000 Plus. Stimulus presentation was controlled by MonkeyLogic. In total, data were collected from 332 MT and 291 V4 neurons across 108 sessions; analyses focused on 15 ensembles including a total of 300 MT and 147 V4 neurons. Each ensemble included neurons from one area with overlapping RFs, and similar ST location and grid positioning. See refs^53,54^ for further details.

### Empirical RF estimation

Neuronal receptive fields were estimated from responses evoked by brief probe stimuli presented during fixation epochs before and after saccades. For each probe position, spike trains were aligned to probe onset and averaged across all repetitions occurring at least 100 ms before or after saccade initiation. Responses were computed in a 0–200 ms post-stimulus window and smoothed using a Gaussian kernel with 5 ms full width at half maximum. These probe-aligned responses provided an estimate of the temporal RF. Spatial RFs were derived by averaging neural activity within each neuron’s response latency window (approximately 50–120 ms) across all probe locations. Presaccadic RF was defined as the probe positions that produced the highest firing rates during fixation before the saccade. For each ensemble, RF area is defined as the center of mass of the RF center of all neurons in an ensemble plus its 8 neighboring probes.

### Encoding model framework

To characterize the fast, time-varying spatiotemporal sensitivity of neurons around saccades, we used an encoding model termed SVGLM, which was previously developed by Niknam et al^53^. This model extends the standard point-process GLM by allowing stimulus sensitivity to vary as a function of time relative to saccade onset while enforcing sparsity over a high-dimensional spatiotemporal parameter space. Neuronal spiking activity was modeled as an inhomogeneous Poisson point process with conditional intensity function (CIF) given by

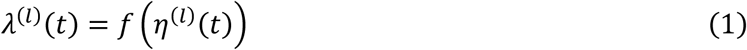

where *λ*^(*l*)^(*t*) is the instantaneous firing rate of the neuron at time *t* in trial *l*, *f*(.) is a sigmoidal nonlinearity, and *η*^(*l*)^(*t*) is the linear predictor capturing the combined effects of sensory input, spike history, and baseline activity defined as

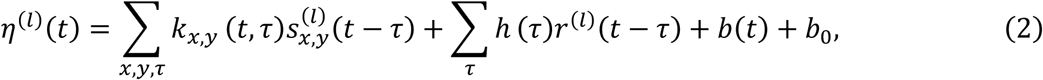

where, 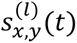 is either 0 or 1 representing respectively the off or on condition in a sequence of probe stimuli presented on the screen at probe location (*x*, *y*) in trial *l* with delay *τ*, evaluated at response time *t*. *r*^(*l*)^(*t*) denotes the spiking response of the neuron for trial *l* and time *t*, *k_x_*_,*y*_(*t*, *τ*) represents the stimulus kernels, ℎ(*τ*) indicates the post-spike kernel applied to the spike history which captures the refractory effect, *b*(*t*) is a time-varying offset kernel capturing baseline firing rate modulations relative to saccade onset. The constant *b*_0_ = *f*^−1^(*r*_0_) with *r*_0_ as the measured mean firing rate (Hz) across all trials in the experimental session, and 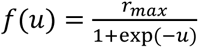 is a static sigmoidal function that describes the nonlinear properties of spike generation with *rm*a*x* indicating the maximum firing rate of the neuron obtained empirically from the experimental data.

To estimate the spatiotemporal sensitivity kernels *k_x_*_,*y*_(*t*, *τ*), space, stimulus delay, and response time relative to saccade onset were discretized into bins, forming a four-dimensional space of STUs. Each STU corresponds to a unique combination of (*x*, *y*, *t*, *τ*), with spatial bins spanning probe locations in a two-dimensional plane and temporal bins (7 ms) spanning both stimulus delay *τ* and response time relative to saccade onset *t*. The sensitivity kernel was expressed as a weighted linear combination of these STUs. Because the full STU space is high-dimensional, sparsity constraints were imposed on the STU weights to isolate the stimulus-response components that significantly contribute to spiking activity. Model parameters were estimated by maximizing a penalized log-likelihood function at the single-trial level^67^,

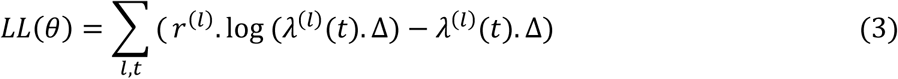

where *θ* denotes the set of parameters used to describe the model kernels, and Δ is the bin size of spike counts. This procedure effectively excluded the majority of STUs, yielding a compact representation of each neuron’s time-varying spatiotemporal sensitivity. Therefore, the resulting model provides a time-resolved description of neuronal sensitivity across space and stimulus history throughout the saccade. More details about the model structure and estimation can be found in ref.^53^.

### Measuring temporal sensitivity

Neurons recorded with the same ST position and probe arrangements (grid positioning and spacing), and with similar RF locations were grouped as an ensemble. Fifteen ensembles were formed, each with a minimum of 10 neurons from one area. Before any analysis, kernels of all neurons were smoothed by moving average using sliding windows of length 50 ms across time *t* and 20 ms across delay *τ* to reduce noise. For each particular time and delay, we constructed a population kernel matrix consisting of the kernel values in the RF area with all neurons in an ensemble. Figure 1c shows nine population kernel matrices at sample delays and times relative to saccade onset for neurons in an example ensemble and probes inside the RF area. To measure the similarity between kernels at different delays and times relative to saccade onset, we vectorized each population kernel matrix and computed the correlation between the kernel vectors at a specific delay and other delay bins at each time to saccade onset for all neurons in an ensemble inside the RF area (Fig. 1f). The correlation was measured for each delay bin and repeated each time to saccade. Thus, at each time to saccade onset, we have a matrix of correlation values each corresponding to the correlation between delays of 0-200 ms binned at 5 ms windows. These correlation maps are shown at some time windows for a sample ensemble in Fig. 2a. Next, we cross-sectioned these correlation maps at each delay bin. Using correlation values obtained from the cross-section of the lower triangle of these correlation maps as sensitivity of each delay bin to other bins, we fitted a sigmoidal function to these values as

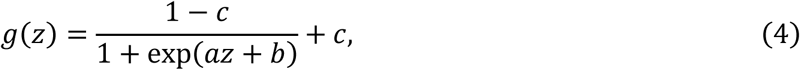

with *a*, *b*, and *c* being the parameters fitted using nonlinear least squares method. The line plot in Fig. 2b shows a sample fit at time -200 ms from saccade onset and delay 20 ms. To measure temporal sensitivity, we defined an index called time difference, which is inversely related to temporal sensitivity. Time difference at each time to saccade onset and delay was defined as the time difference between the measured delay at the inflection point and the actual delay, and it was computed for all 15 ensembles. These time difference values were used to construct time difference maps across time and delay. Figure 2c, d shows an example time difference map for an ensemble and its cross-section at delay 40:60 ms. Before averaging the time difference maps of all 15 ensembles, the original time difference of each ensemble was normalized to range from 0 to 1 using the following formula: (difference−min(difference))/(max(difference)−min(difference)). The average time difference map across 15 ensemble is shown in Fig. 2e. we then averaged this map over delay 40:60ms and corrected the line to have baseline value of zero in fixation period i.e. the mean time difference value in window of -300:-150 ms was subtracted from the average time difference values in delay 40:60 ms (Fig. 2f). We used one-sided Wilcoxon signed-rank test to report p-values for all our statistical comparison analysis, if not mentioned specifically.

### Measuring temporal bias

To measure the similarity between ensemble kernels at each time relative to saccade onset and those in the fixation window, we computed the correlation of the ensemble kernel at a specific delay and time with other delay bins in the fixation period (-500:-300 ms from saccade onset). The correlation was measured for each delay bin and repeated for each time to saccade. Thus, at each time to saccade onset, we have a matrix of correlation values each element corresponding to the correlation between delays of 0-200 ms binned at 5 ms windows of different time windows and those of fixation window. These correlation maps are shown at some time windows for a sample ensemble in Fig. 4c. Next, we measured the similarity between each 300 ms sliding window of these maps by computing the correlation between each map in this window and its central map, meaning that we have a scalar value of correlation associated with each time to saccade onset. The time to saccade corresponding to the central map is considered as the real time and sum of all the time to saccade bins in the sliding window is weighted by their associated correlation value and defined as the perceived time to saccade onset, such that at time t relative to saccade, we compute the perceived time t_perc_ in each 300 ms sliding window as:

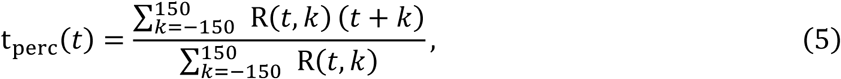

where R(*t*, *k*) is correlation between correlation maps at time t and t+k (correlation map at time t is computed as before by correlating kernels at time t and those at fixation across delays). Figure 4d shows an example plot for real vs perceived time for an ensemble. The average plot of real vs perceived time across 15 ensembles is shown in Fig. 4e.

### Identifying modulated STUs

Modulated STUs are defined as spatiotemporal units where the prevalence of STUs, observed within a 3×3 window surrounding a particular STU’s time and delay, is significantly different during perisaccadic stimulus presentations compared to presentations during stable fixation. To identify STUs whose effect on spiking exhibited systematic temporal modulation, we evaluated changes in the conditional spiking probability *p*(*τ_n_*, *t_m_*) for each delay bin *τ_n_* at time *t_m_*. We compared this quantity to two reference probability profiles, *p*_1_(*τ_n_*) and *p*_2_(*τ_n_*), estimated from non-overlapping temporal baselines. For each (*τ_n_*, *t_m_*) coordinate, temporal modulation was considered significant when the geometric mean of the absolute deviations from both reference profiles exceeded a fixed threshold ℎ^54^:

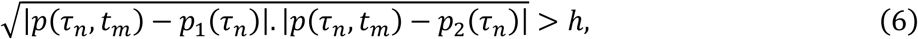

where *p*(*τ_n_*, *t_m_*) is the prevalence of STUs at a given time *t_m_* and delay *τ_n_* within a 3×3 window surrounding it. *p*_1_(*τ_n_*) is the prevalence of STUs during the initial fixation period before the saccade, and *p*_2_(*τ_n_*) is the prevalence of STUs during the fixation period after the saccade. ℎ is a significance threshold, which is set at 0.3.

### Identifying bias-relevant STUs

A subset of modulated STUs were identified as contributing to temporal bias, termed ‘bias-relevant STUs’. The contribution of each modulated STU to the temporal bias was quantified by evaluating its impact on the sensitivity of delay bins across a saccade, by removing each modulated STU one at a time and testing whether the change in kernels across delay is significant based on an index. The squared difference between stimulus kernels of the fitted models at each delay bin (*τ*) and the average kernel over delay of 40:100 ms inside RF area across different times to the saccade (t), was quantified as

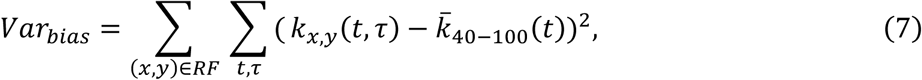

where 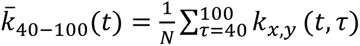 and *N* = 4 is the number of delay bins.

For each neuron within each ensemble, we first computed the variance-based kernel difference metric *Var_sℎift_* using the full model, in which all STU weights were left unaltered. To determine the contribution of individual modulated STUs to this measure, we iteratively nulled STUs one at a time in the model by setting the weight of the selected STU to zero, then recomputed *Var_bias_* after each removal. This yielded a neuron-specific distribution of *Var_bias_* values, each indexed to the absence of a single STU. For each STU, we then defined a temporal bias index as the absolute difference between its perturbed and unperturbed influence on the kernel difference variance:

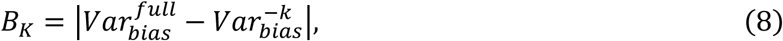

where 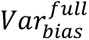 is the index from the full model and 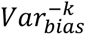 is the index after removing STU *k*. To extract STUs with meaningful impact on temporal mapping, we applied a threshold set to the 90th percentile of the nonzero bias index distribution, corresponding to an index value of 0.56 (Supplementary Figure 3a). STUs satisfying *B_k_* > 0.56 were labeled as temporal bias-relevant, indicating that their removal caused a stronger-than-expected change in kernel weights variance across delay. Figure 6b shows the map of bias-relevant STUs for one representative ensemble of neurons, and Figure 6c shows the mean prevalence map across 15 ensembles of neurons.

### Identifying noise-relevant STUs

Noise-relevant STUs refers to a subset of modulated STUs that contribute to variability across time to saccade. The contribution of each modulated STU to the variability was quantified by evaluating its impact on the sensitivity of time to saccades, by removing each modulated STU one at a time and testing whether the change in kernels across time to saccade is significant based on an index. The standard variance between stimulus kernels of the fitted models across time to saccade (t) at each delay bin (*τ*) inside RF area, was quantified as

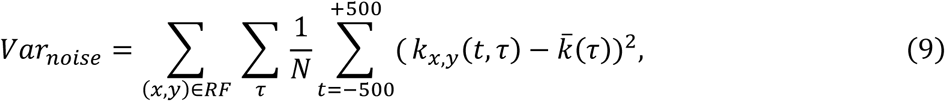

where 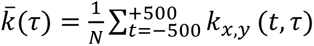 and *N* = 200 is the number of time to saccade bins.

Following the same procedure as the previous section, for each neuron within each ensemble, we computed *Var_noise_* using the full model, and also after removing each modulated STU. For each STU, we then defined a noise index as the absolute difference between its perturbed and unperturbed influence on the kernel weight variance:

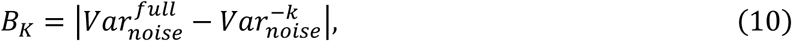

where 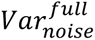 is the index from the full model and 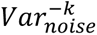 is the index after removing STU *k*. To extract STUs with meaningful impact on temporal mapping, we applied a threshold set to the 90th percentile of the nonzero noise index distribution, corresponding to an index value of 0.05 (Supplementary Figure 3b). STUs satisfying *B_k_*> 0.05 were labeled as noise-relevant, indicating that their removal caused a stronger-than-expected change in kernel weights variance across time to saccade. Figure 7b shows the map of noise-relevant STUs for one representative ensemble of neurons, and Figure 7c shows the mean prevalence map across 15 ensembles of neurons.

## Data and code availability

The datasets generated and/or analyzed for this study will be available on a public repository upon the acceptance of the manuscript.

### Acknowledgements

The authors would like to thank the animal care personnel at the University of Utah. We specifically thank Rochelle D. Moore and Dr. Tyler Davis for their assistance with the NHP experiments. This work was supported by NIH EY026924 to B.N.; NIH EY031477 to N.N.; AFOSR FA9550-24-1-0235 to N.N. and B.N.; NIH EY014800 and an Unrestricted Grant from Research to Prevent Blindness, New York, NY, to the Department of Ophthalmology & Visual Sciences, University of Utah.

## Author contributions

N.N. and B.N. conceived the study. B.N. performed the surgical procedures. B.N., N.N., and A.A. designed the experiment. B.N., N.N., K.C., M.Z., A.P., and G.W. designed the analysis. B.N. and A.A. performed the physiology experiments and acquired data. A.A. performed the modeling. A.P. performed the data analysis. A.P., K.C., N.N., B.N., M.Z., and H.R. wrote the manuscript.

## Competing interests

The authors declare no competing interests.

**Supplementary Information** is available for this paper.

