## Supplementary Information for "A population readout of extrastriate activity reveals biased and smoothed temporal representations across saccades"

Neda Nategh

Behrad Noudoost

Maryam Zekri

**Word counts**

833

**Number of figures**

4

### Dimensionality reduction

The rapid perisaccadic modulation of neuronal spatial sensitivity necessitates a high-dimensional representation of spatiotemporal response properties. At any time relative to saccade onset, neuronal responses depend on the spatial location of the stimulus and on the delay between stimulus presentation and spike generation.

To make the problem tractable, we first discretized both the ~1 s of response time relative to saccade onset (t) and the 200 ms of stimulus-response delay ($\tau$) into 7-ms bins, matching the stimulus update rate. Although this step reduced dimensionality substantially (~$10^7$), the resulting spatiotemporal space remained too large for reliable estimation with the available data. We therefore introduced a parametric representation of temporal structure using smooth basis functions. Specifically, we modeled the temporal dependence of neuronal sensitivity using separable basis functions defined as

$$\psi_{i,j}(t,\tau) = \mathcal{U}_i(\tau)\,\mathcal{V}_j(t) \tag{1}$$

where $\mathcal{U}_i(\tau)$ and $\mathcal{V}_j(t)$ are second-order B-spline functions spanning the delay and time dimensions, respectively. The delay basis $\mathcal{U}_i(\tau)$ covered a 200-ms window using 33 uniformly spaced knots at {-13, 6, …, 204, 211} ms, yielding 30 basis functions. The time basis $\mathcal{V}_j(t)$ spanned a 1081-ms interval centered on saccade onset, defined by 159 uniformly spaced knots at {-554, -547, …, 545, 552} ms and yielding 156 basis functions. Knot spacing followed the original 7-ms temporal discretization, ensuring that sensitivity could vary at the same temporal resolution as the stimulus. This basis expansion reduced the dimensionality of the spatiotemporal kernel representation by approximately two orders of magnitude.

Even with this reduction, the number of spatiotemporal units (STUs) remained too large for stable estimation given the limited number of spikes available during the brief perisaccadic period. To further constrain the model, we applied a data-driven pruning procedure to identify STUs that made a statistically meaningful contribution to neuronal firing[1]. For each STU, we assessed its significance by fitting a simplified generalized linear model (GLM)[2–4] in which that STU was evaluated independently. Model fitting was repeated on 100 randomly selected subsets of spike trains (each containing 35% of trials) to obtain a distribution of weight estimates. A control distribution was generated by fitting the same model to 100 subsets of shuffled spike trains, thereby disrupting the stimulus-response relationship. The conditional intensity function (CIF) of this GLM was defined as[5,6]

$$\lambda_t = f\left(\sum_{\tau=1}^{T} s_{x,y}\,(t-\tau)\,\kappa_{x,y,i,j}\,\psi_{i,j}(t,\tau)\right) \tag{2}$$

where $\lambda_t$ denotes the instantaneous firing rate, $s_{x,y}$ is the stimulus history of length $T$ at spatial location $(x, y)$, $\kappa_{x,y,i,j}$ is the weight associated with the STU at location $(x, y)$, and time and delay bin indices $(i, j)$, and $f(.)$ is a static nonlinearity. An STU is discarded if:

$$|\mu - \tilde{\mu}| \geq 1.5\,\tilde{\sigma} \tag{3}$$

where $\mu$ is the mean weight from the original data, and $\tilde{\mu}$ and $\tilde{\sigma}$ are the mean and standard deviation of the shuffled control distribution. The threshold of 1.5 was chosen heuristically to balance dimensionality reduction with retention of relevant stimulus features. Applying this criterion reduced the number of STUs to approximately $10^4$, yielding a parameter space suitable for robust model fitting without overfitting. Finally, we fit the sparse variable generalized linear model (SVGLM) using only this subset of significant STUs in the linear filtering stage of the model according to the CIF defined in Eq. (1) and (2) in Methods. The spatiotemporal sensitivity kernel for each spatial location was expressed as a weighted sum of the retained basis functions,

$$k_{x,y}(t,\tau) = \sum_{i,j} \kappa_{x,y,i,j}\;\psi_{i,j}(t,\tau) \tag{4}$$

with weights corresponding to non-significant STUs fixed at zero. This low-dimensional representation enabled reliable estimation of time-varying spatiotemporal sensitivity maps and allowed us to characterize how neuronal encoding properties evolve across the course of a saccadic eye movement.

**Model performance**

Model performance was evaluated exclusively on held-out test data, with the dataset split into training (35%), validation (30%), and testing (35%) subsets. Supplementary Figure 1a shows single-trial examples, where the good and average trials were selected based on the largest and median normalized log-likelihood difference ($\Delta LL$ per spike), respectively. Supplementary Figure 1b summarizes model performance using $\Delta LL$ per spike, defined as the log-likelihood of the observed spike train under the model-predicted firing rate minus that under a null model with constant mean firing rate, normalized by the total spike count. Comparing the perisaccadic window (0–150 ms after saccade onset) with the fixation window (-300 to -150 ms) revealed slightly higher information during the perisaccadic period (0.16±0.006 vs. 0.15±0.005 bits/sp, Wilcoxon signed-rank $p=3.48\times10^{-5}$), indicating that the model accurately tracks spike timing even during rapid changes in neural sensitivity. Supplementary Figure 1c then reports firing-rate prediction accuracy, quantified by the correlation coefficient (CC) between the model-predicted and empirical responses to repeated 300-ms probe sequences during fixation (-500 to -200 ms relative to saccade onset). Spike trains were binned in non-overlapping 30-ms windows, smoothed with a 5 ms full-width-at-half-maximum Gaussian kernel, and normalized to zero mean and unit variance. Intrinsic data reliability was estimated from 15 repetitions of random 60%–40% splits of the neural responses to compute the data–data CC. The data–data CC (0.44±0.01) was significantly higher than the model–data CC (0.30±0.01, $p=5.96\times10^{-75}$), showing that while neural variability limits predictability, the model captures a substantial fraction of the explainable firing-rate structure.

### Supplementary figures and figure legends

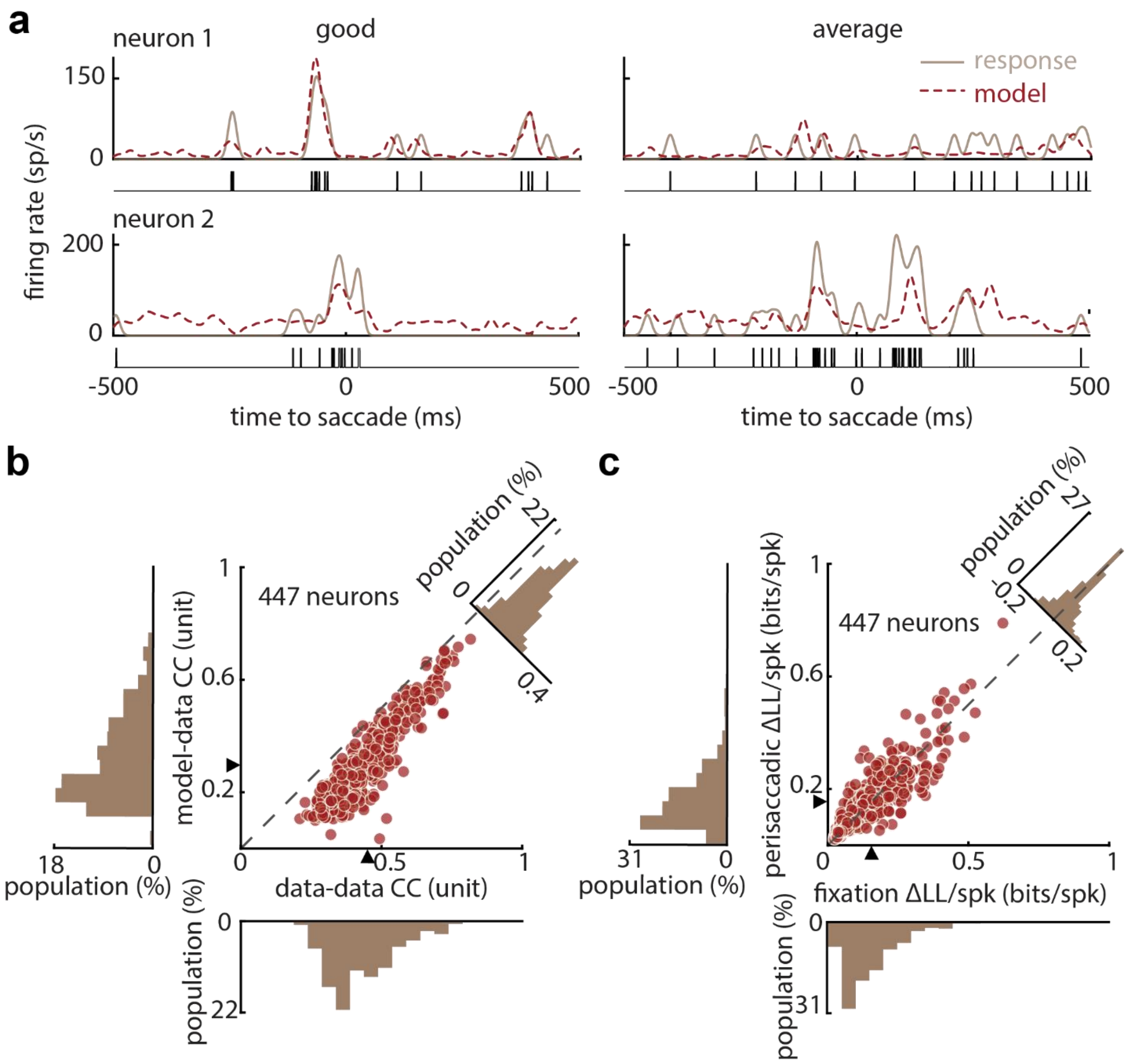


**Supplementary Fig. 1: Model performance in predicting the firing rates. a.** Example single-trial responses for two neurons illustrating model predictions at the level of spike timing. For each neuron, a 'good' trial (highest normalized $\Delta LL$ per spike) and an 'average' trial (median $\Delta LL$ per spike) are shown. Solid traces indicate the smoothed empirical firing rate, dashed traces show the model-predicted firing rate, and vertical ticks denote individual spikes (empirical data). **b.**

Firing-rate prediction accuracy quantified by the correlation coefficient (CC) between model-predicted and empirical responses to repeated 300-ms probe sequences during fixation ($-500$ to $-200$ ms relative to saccade onset). Each point corresponds to one neuron. The intrinsic reliability of the data (data–data CC) was estimated from 15 random 60%–40% splits of the neural responses and was significantly higher than the model–data CC (data–data =0.44±0.01, median=0.45, model–data =0.30±0.01, median=0.30; $p$=5.96×$10^{-75}$). Marginal histograms show the distributions of data–data and model–data CCs, and the upper-right panel shows the distribution of their differences. **c.** Population comparison of normalized log-likelihood improvement ($\Delta LL$ per spike) between fixation (-300 to -150 ms) and the perisaccadic interval (0 to 150 ms). Histograms display the marginal distributions for each condition (left and bottom) and their pairwise differences (upper right). The medians are indicated by black triangles (fixation median=0.15 bits/spike; perisaccadic median=0.16 bits/spike). The median information conveyed by single spikes was slightly higher in the perisaccadic period than during fixation (fixation =0.15±0.005 bits/spike, perisaccadic =0.16±0.006 bits/spike; Wilcoxon signed-rank test $p$=3.48×$10^{-5}$), indicating improved model fit during the perisaccadic period.

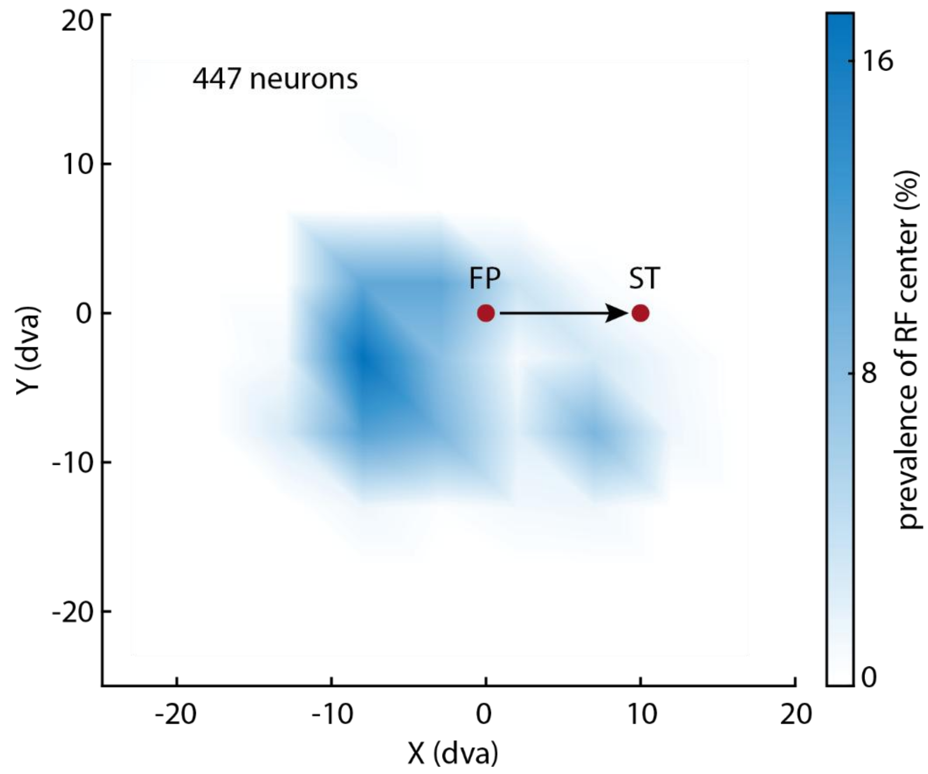


**Supplementary Fig. 2: Spatial distribution of all neuronal RF centers relative to the FP and ST.** Recordings from ensembles recorded with leftward saccades were mirrored to combine with rightward saccade data. The color bar indicates the percentage of neurons whose RF centers fall within each location.

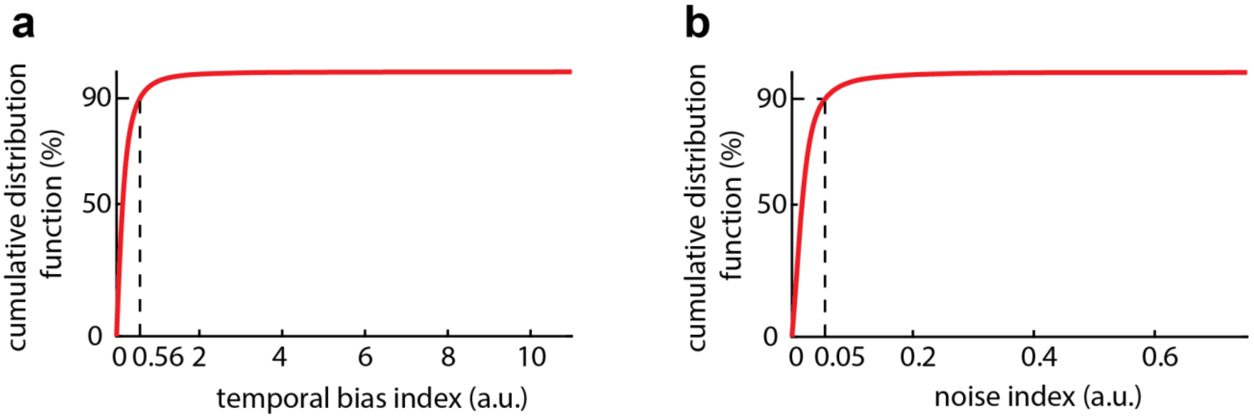


**Supplementary Fig. 3 Identifying bias- and noise-relevant STUs using indices.** The cumulative distribution function of all the non-zero **a.** temporal bias and **b.** noise indices. Using the 90th percentile as a threshold, the STUs with an absolute bias index difference above 0.56 are defined as bias-relevant and those with an absolute noise index difference above 0.05 are identified as noise-relevant.

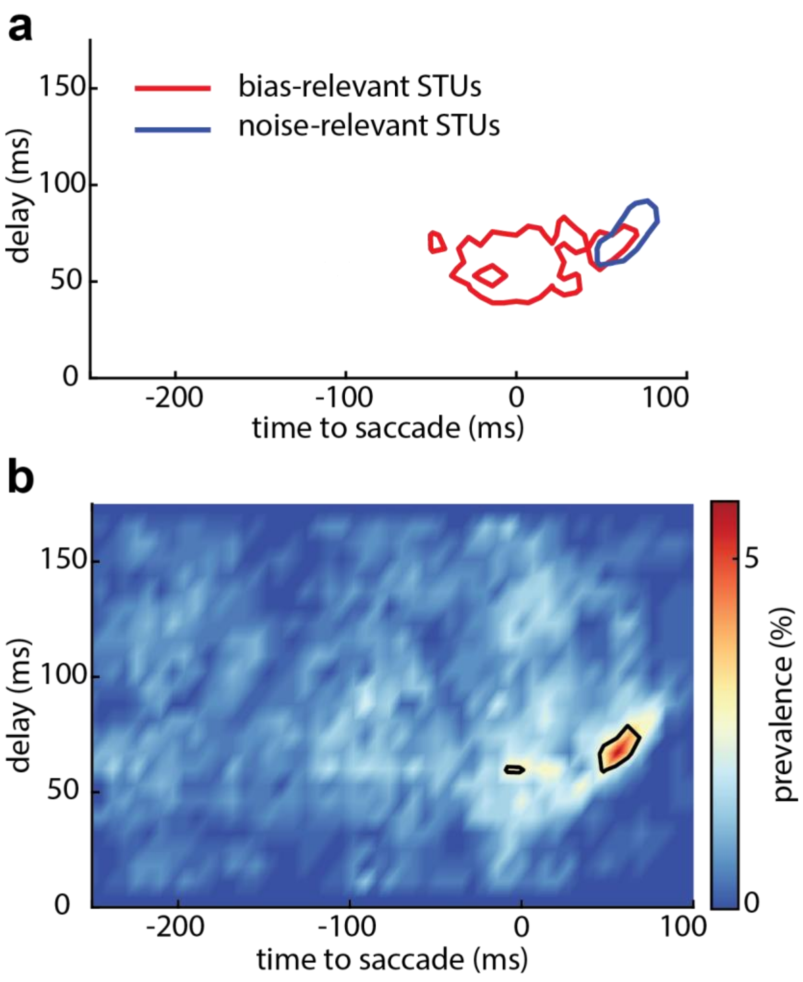


**Supplementary Fig. 4: The overlapping between bias- and noise- relevant STUs. a.** Red line illustrates the contour of bias-relevant STU prevalence above 50% of the peak prevalence. Blue line is the same contour for noise-relevant STUs. **b.** The prevalence map for overlapping STUs that are both bias- and noise-relevant.
